# Early-life density-dependent mortality in the Gulf of Thailand

**DOI:** 10.64898/2026.02.17.706505

**Authors:** Jeeratorn Yuttharax, Pavarot Noranarttragoon, Changoma Marko Fransis, Methee Kaewnern, Matsuishi Takashi Fritz

## Abstract

Density-dependent mortality is a key mechanism regulating fish population dynamics, yet empirical evidence from tropical systems remains limited. This study investigated density-dependent survival during early life stages of 25 dominant taxa from trawl landings in the Gulf of Thailand, a productive tropical system dominated by multispecies and multi-gear fisheries, where trash fish (TF) catches from trawlers comprise proportions of juveniles. Temporal variation in the abundance of fish at ages smaller than *L*_25_ was linked to subsequent survival patterns to evaluate compensatory dynamics. Growth parameters revealed wide interspecific variation in life-history strategies, ranging from fast-growing (*Encrasicholina* spp.) to a slower-growing demersal group (Sciaenidae). Most taxa exhibited a negative density– survival relationship, with 11 taxa (including *Rastrelliger kanagurta, Atule mate*, and *Lutjanus lutjanus*) showing strong and significant compensatory density dependence (*p* < 0.001), indicating elevated mortality at higher juvenile densities. Evidence for depensatory dynamics was weak and statistically unsupported. These results provide empirical biological evidence that compensatory density-dependent mortality commonly operates during early life stages in tropical multispecies and multi-gear fisheries. The findings suggest that juvenile removal via TF catches does not uniformly translate into reduced long-term productivity, highlighting the importance of incorporating early-life density dependence into stock assessment and management frameworks.

## Introduction

Density-dependent mortality is a fundamental process shaping population dynamics in fishes, particularly during early juvenile stages when individuals might transition from pelagic to demersal habitats and competition for space and resources intensifies. In fisheries science, such processes are commonly conceptualized as compensatory mechanisms, whereby increased population density often leads to higher mortality but can function to stabilize survival and recruitment across cohorts (Myers and Cadigan 1993; Lorenzen 2000; Rose et al. 2001; Andersen et al. 2017; Stige et al. 2019). Under strong compensatory dynamics, reductions in juvenile abundance may have limited effects on subsequent recruitment, as density-dependent processes buffer population fluctuations (Hazlerigg et al. 2012).

Accordingly, fisheries models typically embed compensatory density dependence through stock–recruitment relationships linking spawning stock biomass to the number of new recruits (Ricker 1954; Beverton and Holt 1957; Myers and Cadigan 1993). Empirical support for this assumption is substantial for many fish populations, particularly in temperate systems (Ohlberger et al. 2014; Andersen et al. 2017; Lorenzen and Camp 2018; Zimmermann et al. 2018; Stige et al. 2019). By contrast, direct biological evidence from tropical marine ecosystems remains limited and is largely derived from studies that were conducted in coral reef systems (Jenkins et al. 1991; Doherty et al. 2004; Houk et al. 2018, 2020; Taylor et al. 2019). High species diversity, overlapping cohorts, prolonged or year-round spawning, and multi-gear exploitation obscure stock–recruitment signals in tropical fisheries, complicating inference on the timing, strength, and mechanisms of density dependence (Pauly 1980; Longhurst and Pauly 1987; Hilborn and Walters 1992). Consequently, compensatory density dependence, despite its central role in fisheries theory, remains poorly resolved for tropical systems.

In tropical multispecies fisheries, much of the bycatch that would be discarded in temperate fisheries is retained and utilized as trash fish (TF), particularly in trawl fisheries. TF predominantly consists of juvenile individuals that are smaller than the size at recruitment to the fishery, defined here as the size at which fish first become vulnerable to fishing gears. However, species-specific growth trajectories mean that TF may also include individuals that fall within post-recruitment size classes (Staples and Funge-Smith 2005; Pérez Roda et al. 2019). Concerns surrounding the utilization of juveniles in TF catches have persisted for decades, yet a critical uncertainty remains: whether the removal of early life stages of potentially commercial species in TF catches necessarily translates into reduced long-term availability of high-quality fish for human consumption (Christensen 1996; Pauly et al. 1998; Staples and Funge-Smith 2005; FAO 2022). Resolving this question requires explicit consideration of density-dependent survival during early life stages.

Measures designed to reduce bycatch and TF, such as gear modifications, changes in fishing practices, spatial or temporal restrictions, and discard bans, are directly linked to the volume and composition of landed catches (Kelleher 2005; Suuronen and Sardà 2007; FAO 2011; Pérez Roda et al. 2019). Gear-based measures often implicitly assume that escaped or released fish survive and later contribute to recruitment. However, post-escape survival is frequently either low or uncertain, particularly when release occurs late in the capture process (Main and Sangster 1990; Suuronen 2005; Broadhurst et al. 2006; Gilman et al. 2013; He 2015). In non-selective trawl fisheries, fish are harvested across multiple life stages that differ in ecological requirements and vulnerability to density-dependent processes. This highlights the need for empirical evidence on early-life abundance and survival to better understand recruitment dynamics in tropical multispecies systems (Rose et al. 2001; Stige et al. 2019).

The Gulf of Thailand (GoT) is a key region for global fishmeal production, which relies largely on TF, mainly originating from trawlers, as the primary raw material (Funge-Smith et al. 2005; Staples and Funge-Smith 2005, Yuttharax et al. 2026). This semi-enclosed gulf is a dynamic tropical marine ecosystem bordered by Thailand, Cambodia, Vietnam, and Malaysia. It encompasses a surface area of approximately 320,000 km^2^, with a mean depth of 45 m and a maximum depth of 80 m (Sherman and Hempel 2009). The GoT is characterized by high primary productivity and supports complex multispecies, multi-gear fisheries where diverse species across multiple trophic levels are harvested simultaneously. This complexity presents significant challenges to sustainable management (Pope 1979; FAO 2023; Matsuishi 2025).

Despite the ecological and economic importance of TF as a whole, data on populations of the species comprising TF catches remain critically understudied, as fisheries research has historically prioritized high-value target species for human consumption. While the GoT is a shared marine ecosystem bordered by several countries, comprehensive long-term data on TF populations are available only from Thailand, where systematic trawl fisheries surveys and national landing statistics have been consistently collected, including detailed species composition and length-frequency information for TF catches (Kulanujaree et al. 2020; Noranarttragoon et al. 2023; Yuttharax et al. 2026). This unique data availability reflects Thailand’s long-standing monitoring of trawl fisheries and TF utilization, providing a rare opportunity to examine early-life abundance patterns across multiple taxa and to explore density-dependent survival processes in a tropical multispecies fishery.

This study integrates monthly commercial fisheries survey data with annual landing statistics to investigate density-dependent survival during early life stages of fish populations in the GoT. By linking temporal variation in the abundance of young fish (defined by sizes smaller than the length at which 25% of consumable fish were captured [*L*_25_]) to subsequent survival patterns, the study provides empirical biological evidence on the strength and timing of density-dependent mortality in a tropical multispecies and multi-gear fishery system.

## Materials and methods

### Data sources and preparation

This study utilized trawl fisheries data as an effective observational source for early-life population dynamics, as they capture high abundances of juvenile and small -sized individuals across multiple species. Two datasets of trawl fisheries in the GoT, covering the period 2016–2023, same study area described in Yuttharax et al. (2026) (Fig. 1), were obtained from Thailand’s Department of Fisheries as part of the country’s routine national fisheries monitoring programs, consisting of catch statistics and survey time-series data.

**Fig. 1:**
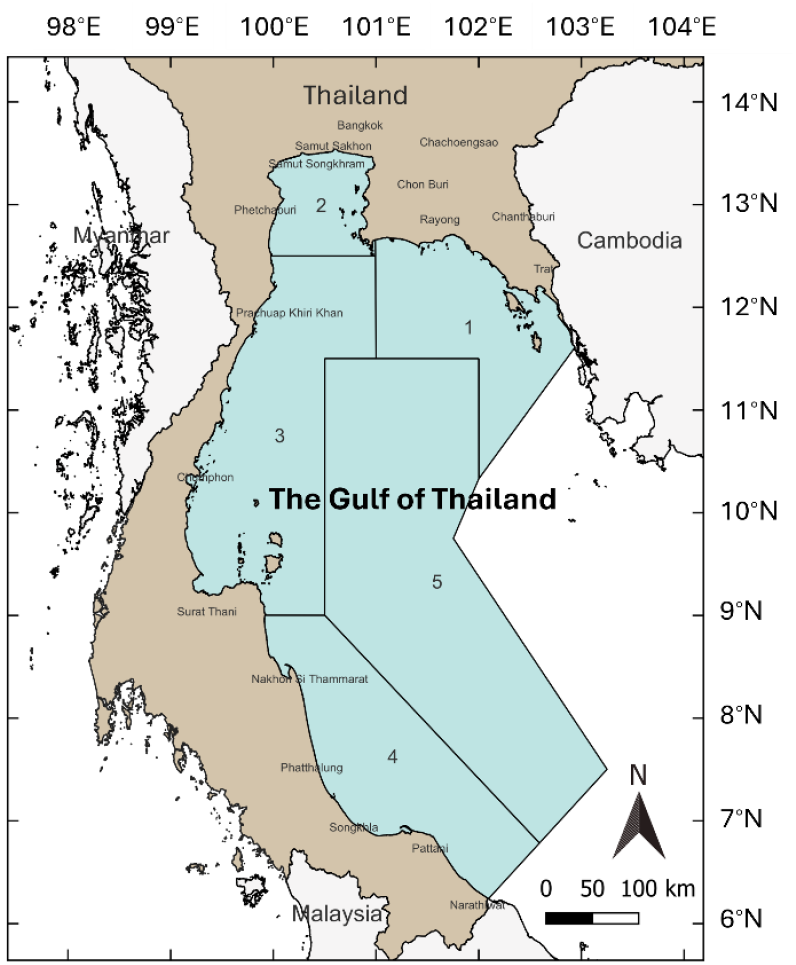
Geographic location of the study area in the Gulf of Thailand with fisheries statistical zones, reproduced from Yuttharax et al. (2026)

Catch statistics are compiled annually by the Fisheries Statistics Group, Fisheries Development Policy and Planning Division, and have been publicly available online since 2009 (https://www4.fisheries.go.th/local/index.php/main/site/strategy-stat [accessed 23 December 2024]). Survey data were originally collected and recorded by five regional centers of the Marine Fisheries Research and Development Division and then compiled by the Fisheries Resource Assessment Group. Those data were obtained at landing sites in Thailand along the GoT coast once a month.

The total trawl landings sampled during 2016–2023 was 4,015 at the trip-level: 2,514 sampled trips for otter-board trawl (OBT), 1,219 for pair trawl (PT), and 282 for beam trawl (BT). Species were identified by well-trained fisheries scientists following the taxonomic guides of Carpenter and Niem (1998a, b; 1999a, b; 2001a, b) and were categorized either as consumable fish (CF) intended for human consumption or as TF destined for non-food uses. For each economically important species, up to 100 individuals were sampled from each landing. All individuals were counted and then collectively weighed in grams, rather than weighed individually. All length measurements were recorded to the nearest 0.5 cm. Fish were measured as total length, except for *Megalaspis cordyla*, for which fork length was used, while squid and cuttlefish were measured by mantle length. Sex determination was not conducted.

We conducted comprehensive data cleaning and validation to correct entry errors and remove inconsistencies. Species-level catch weights (kg) from the trawl surveys were then raised to annual catch totals (tonnes) using proportional contributions derived from annual statistics collected by the Department of Fisheries. The top 25 taxa with the highest catch volumes in total trawl catches from the raised-catch data, representing 39.82% of the total landings from trawlers, and for which length-frequency (lfq) data have been collected consistently, were selected to observe density-dependent mechanisms.

Trophic-level estimates for each taxon were used in conjunction with the study results to interpret their role within the marine ecosystem. Trophic levels for fish species were compiled from FishBase (www.fishbase.se), while those for invertebrates were obtained from SeaLifeBase (www.sealifebase.se). For taxa not recorded to species level, or where a species-specific trophic-level estimate was not available, a trophic level was assigned by averaging the trophic levels of all species in the most closely related taxonomic group (i.e. within a higher-level taxon like genus, family, or order) (Yuttharax et al. 2026).

### Length-frequency distributions and selectivity

The lfq distributions of 25 taxa were analyzed using monthly lfq data. Only OBT and PT data were combined, as BT contributed minimally to trawl catches (especially that of TF at 0.20% of TF landings in the GoT) and the sampling weight and lfq data for that gear were limited (Yuttharax et al. 2026). Moreover, to remove biologically unreliable lfq data, samples with the lowest sampling ratios for each taxon and landing, which together accounted for <5% of the total fish frequency, were excluded from the analysis. For each month, the number of individuals in landing *N*_*l*_ in each length class *i* in month *m* was calculated as:

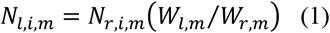

where *N*_*r,i,m*_ is the number of individuals in the length class *i* randomly sampled in month *m*; *W*_*l,m*_ is the total landing weight in month *m*; and *W*_*r,m*_ is the total weight of individuals randomly sampled and measured for length in month *m*. Through this procedure, the lfq distribution obtained from the sample was adjusted to represent the species’ monthly landings.

The selectivity of CF catches was analyzed to define the pre-recruit stage of each taxon for density-dependent survival analysis, as shown in Table 1. The CF ratios *R* were calculated using the equation:

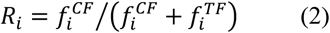

where 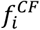 and 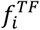 are the numbers of fish in catch groups CF and TF, respectively, in length class *i*. Length class *i* refers to intervals of the recorded fish lengths, and each interval is defined to include fish with lengths *l* within a specific range. These data were then fitted to a logistic regression:

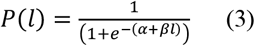

where *P*(*l*) is the probability of capture at length *l*, and *α* and *β* are parameters of the logistic function. The logistic function calculates the probability of a fish taxon being caught at different lengths. *L*_50_ is the length at which 50% of CF are captured, calculated as:

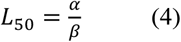

**Table 1.** Length distribution of the top 25 taxa in trawl catches in the Gulf of Thailand during the period 2016–2023.

| No. | Scientific name | CF catch (cm) | TF catch (cm) | Total catch (cm) | $L_{25}$ (cm) | $L_{50}$ (cm) | $L_{75}$ (cm) |
| --- | --- | --- | --- | --- | --- | --- | --- |
| 1 | *Uroteuthis duvaucelii* | 1.5–40.5 | 1.5–14.0 | 1.5–40.5 | 4.24 | 5.10 | 5.95 |
| 2 | *Selaroides leptolepis* | 3.0–23.0 | 1.5–15.0 | 1.5–23.0 | 11.39 | 12.37 | 13.36 |
| 3 | *Sardinella gibbosa* | 6.0–30.0 | 2.0–18.0 | 2.0–30.0 | 11.99 | 14.60 | 17.22 |
| 4 | *Priacanthus tayenus* | 1.0–31.5 | 0.5–20.5 | 0.5–31.5 | 11.04 | 12.27 | 13.51 |
| 5 | *Saurida elongata* | 1.0–61.0 | 5.0–30.0 | 1.0–61.0 | 10.92 | 12.56 | 14.20 |
| 6 | *Scolopsis taenioptera* | 5.0–29.5 | 3.0–20.5 | 3.0–29.5 | 8.86 | 9.90 | 10.95 |
| 7 | Sciaenidae | 3.5–48.0 | 3.0–18.5 | 3.0–48.0 | 11.16 | 13.24 | 15.31 |
| 8 | *Rastrelliger kanagurta* | 2.0–33.5 | 3.5–17.0 | 2.0–33.5 | 13.67 | 14.64 | 15.61 |
| 9 | *Sepia aculeata* | 1.5–29.5 | 1.5–14.0 | 1.5–29.5 | 2.26 | 3.69 | 5.13 |
| 10 | *Rastrelliger brachysoma* | 4.5–26.5 | 3.5–19.0 | 3.5–26.5 | 9.63 | 10.88 | 12.14 |
| 11 | *Encrasicholina* spp. | 9.0–10.5 | 1.0–13.0 | 1.0–13.0 | 10.27 | 10.84 | 11.41 |
| 12 | *Upeneus sulphureus* | 6.0–19.0 | 3.0–14.5 | 3.0–19.0 | 8.91 | 10.06 | 11.21 |
| 13 | Platycephalidae | 2.0–49.0 | 2.5–46.0 | 2.0–49.0 | 11.11 | 13.14 | 15.18 |
| 14 | *Saurida undosquamis* | 2.0–53.5 | 3.0–25.5 | 2.0–53.5 | 12.89 | 14.56 | 16.23 |
| 15 | *Nemipterus mesoprion* | 2.0–24.5 | 2.5–16.5 | 2.0–24.5 | 8.47 | 9.46 | 10.45 |
| 16 | *Nemipterus peronii* | 0.5–29.0 | 2.0–19.0 | 0.5–29.0 | 11.38 | 12.67 | 13.96 |
| 17 | *Nemipterus japonicus* | 1.0–28.5 | 0.5–15.5 | 1.0–28.5 | 8.68 | 9.76 | 10.85 |
| 18 | *Saurida isarankurai* | 7.0–24.5 | 0.5–15.0 | 0.5–24.5 | 11.22 | 12.62 | 14.03 |
| 19 | *Atule mate* | 4.0–49.5 | 3.0–15.0 | 3.0–49.5 | 11.16 | 12.23 | 13.31 |
| 20 | *Lagocephalus spadiceus* | 6.0–33.5 | 4.0–19.0 | 4.0–33.5 | 10.48 | 11.90 | 13.31 |
| 21 | *Siganus canaliculatus* | 7.5–27.5 | 1.0–20.5 | 1.0–27.5 | 15.31 | 17.57 | 19.83 |
| 22 | *Sphyræna jello* | 2.0–135.0 | 4.0–45.0 | 2.0–135.0 | 18.14 | 19.54 | 20.95 |
| 23 | *Megalaspis cordyla* | 4.0–37.5 | 2.0–19.0 | 2.0–37.5 | 13.92 | 15.74 | 17.56 |
| 24 | *Parastromateus niger* | 6.0–59.0 | 4.0–13.0 | 4.0–59.0 | 8.86 | 10.25 | 11.64 |
| 25 | *Lutjanus lutjanus* | 4.0–23.0 | 3.0–18.5 | 3.0–23.0 | 13.95 | 15.26 | 16.57 |

Taxa were categorized as consumable fish (CF) or trash fish (TF) and in relation to size selectivity, where $L_{25}$ , $L_{50}$ , and $L_{75}$ represent the lengths at which 25%, 50%, and 75% of the individuals entering the gear are retained. All fish taxa were measured as total length, except *Megalaspis cordyla* was measured as fork length; squid and cuttlefish were measured by mantle length

The probability of being caught at other lengths provides further detail on the gear selectivity range, calculated with the formula:

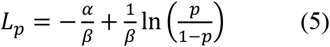

where *p* = 0.25 for *L*_25_, and *p* = 0.75 for *L*_75_ (Sparre and Venema 1998).

### Growth estimation and age–frequency conversion

To quantify density-dependent attributes, age-structured frequency distributions were required for each taxon. Lfq observations were therefore transformed into relative age bins based on species-specific growth trajectories. Growth in length was modelled using the von Bertalanffy growth function (VBGF) (von Bertalanffy 1938):

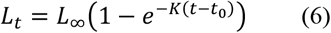

where *L*_∞_ denotes asymptotic length, *K* is the growth coefficient, and *t*_0_ represents the theoretical age at zero length.

Growth parameters were estimated using the ELEFAN_SA routine (Electronic LEngth Frequency ANalysis with simulated annealing) implemented in the TropFishR package (Mildenberger et al. 2017) in R Studio. The lfq dataset was restructured using a moving-average smoothing filter for window sizes of 5 and 7. Starting values for parameter optimization were informed by plausible *K* (growth coefficient) ranges reported in FishBase for fish taxa and SeaLifeBase for invertebrate taxa. For taxa at the family or group level, the bounds of *K* ranges were broadened to encompass growth parameters recorded for GoT species. The third quartile (75th percentile) of the total length data within each taxon was adopted as the initial estimate of *L*_∞_, with a tuning interval of −5 cm to +10 cm around this value. The seasonal growth oscillations function was disabled in the ELEFAN configuration.

Parameter estimation followed a bootstrap resampling design with 100 replicates (Schwamborn et al. 2019; Taufani and Matsuishi 2025a, b). Each run was initialized with a unique random seed to ensure reproducibility and parameter variability. The simulated annealing optimization time was fixed (SA_time = 30) to balance computational efficiency with search depth. From each bootstrap replicate, the estimated values of *L*_∞_, *K*, and the growth performance index: *ϕ*^′^ = log_10_ *K* + 2 log_10_ *L*_∞_ (Pauly and Munro 1984; Schwamborn et al. 2023) were extracted. The median values across bootstrap replicates were adopted as robust estimator values, minimizing sensitivity to outliers. The ELEFAN goodness-of-fit index (*R*_*n*_) was used to evaluate the model performance, and summary statistics (mean, standard deviation, 95% confidence interval) were computed for all parameters.

The estimated VBGF curves were then used to convert lengths into relative monthly ages. The lfq data were projected onto age space using the bootstrap model with the highest median *R*_*n*_ (best-supported growth trajectory). Counts were subsequently re-aggregated into monthly age bins, producing taxon-specific age-frequency matrices used as input for the density-dependent survival analysis.

### Density-dependent survival analysis

Density-dependent mortality is widely recognized as a fundamental mechanism regulating survival during early life stages in marine fishes, particularly prior to recruitment into the exploited population (Andersen et al. 2017). In this study, the pre-recruit stage was operationally defined as individuals smaller than *L*_25_ as observed in the CF catches. This threshold represents the size domain where TF are predominantly retained and therefore provides a biologically meaningful boundary delimiting the phase during which density-dependent processes are expected to exert the strongest influence.

For each of the top 25 taxa occurring in trawl catches in the GoT, age-specific abundance (frequency) was reconstructed from monthly *j* lfq data using the estimated growth curves. For each taxon *i*, the survival rate *S* from age *t* to *t* + 1 was calculated as:

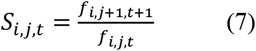

Only *S* values within the biologically meaningful range of 0–1 were retained for the density-dependent analyses. Values outside this interval were treated as artefacts rather than true demographic responses, as they likely reflect sampling noise or disproportionate cohort representation, even after prior data filtering. Excluding these outliers ensured that the analysis captured realistic population dynamics and prevented spurious correlations driven by measurement error rather than biological processes. To quantify density dependence in pre-catch-stage survival, a linear model was assumed:

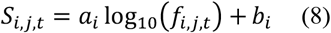

where *a*_*i*_ is the density-dependent slope and *b*_*i*_ is the *y*-intercept, and log_10_(*f*_*i,j,t*_) is the base-10 logarithm of the age-specific frequency for the taxon *i* in month *j* at age *t*. A significantly negative slope (*a*_*i*_ < 0) indicates compensatory density-dependent survival, consistent with well-established mechanisms in early stages of marine fishes (Myers and Cadigan 1993; Lorenzen 2000; Rose et al. 2001; Andersen et al. 2017; Stige et al. 2019).

## Results

### Growth parameters

Growth parameter estimates revealed pronounced interspecific variability among the 25 dominant taxa in trawl catches of the GoT (Table 2) during the period 2016–2023. Asymptotic length *L*_∞_ ranged from 8.0 cm in *Encrasicholina* spp. to 55.2 cm in *Sphyraena jello*, reflecting broad contrasts in life-history strategies. Growth coefficients (*K*) exhibited similarly wide dispersion, from 0.134 year^−1^ (Sciaenidae) to 1.356 year^−1^ (*Encrasicholina* spp.), indicating rapid turnover in small pelagics relative to slower-growing demersal taxa. The growth performance index (*ϕ*^′^) generally increased with body size, with the highest values observed for *Sphyraena jello* and *Atule mate*, suggesting elevated productivity potential among larger predatory fishes.

**Table 2.**
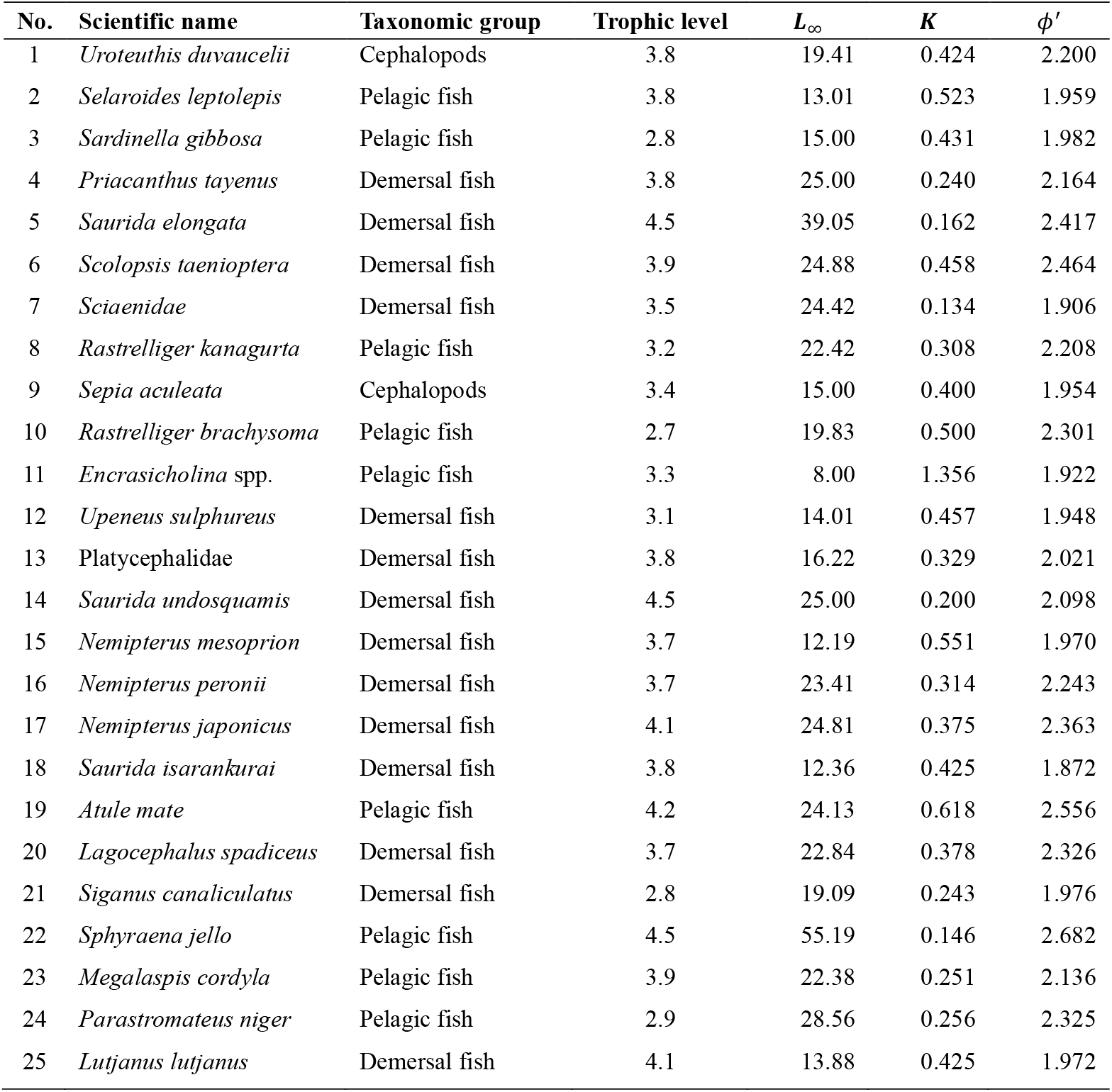
Growth parameters of the top 25 taxa originating from trawl fisheries in the Gulf of Thailand in 2016–2023. *L*_∞_ is asymptotic length, *K* is the growth coefficient, and *ϕ*^′^ is the growth performance index

### Density-dependent mortality

Results of the density-dependent survival analyses revealed marked interspecific variation among the 25 dominant taxa (Table 3). The majority showed negative density-survival slopes, indicating compensatory mortality during early juvenile stages as sizes below *L*_25_, with 11 taxa exhibiting strong support for this mechanism (*p* < 0.001). Species with faster growth or higher ecological turnover (Table 2), such as *Rastrelliger kanagurta, Atule mate*, and *Lutjanus lutjanus*, displayed the clearest compensatory responses, whereas several slow-growing demersal taxa showed weaker signals. Only four taxa demonstrated positive slopes, but none were statistically significant, suggesting that depensatory effects, if present, were weak or masked by sampling variability.

**Table 3.** Density-dependent attributes of the top 25 taxa from trawl fisheries in the Gulf of Thailand from 2016 to 2023, analyzed within ages corresponding to lengths below the length at which 25% of consumable fish are captured (*L*_25_)

| No. | Scientific name | *a* | *b* | $R^2$ | Age < $L_{25}$ |
| --- | --- | --- | --- | --- | --- |
| 1 | *Uroteuthis duvaucelii* | 0.0229 | 0.0082 | 0.020 | +, n.s. |
| 2 | *Selaroides leptolepis* | -0.1246 | 0.9186 | 0.102 | -, *** |
| 3 | *Sardinella gibbosa* | -0.0166 | 0.3605 | 0.002 | -, n.s. |
| 4 | *Priacanthus tayenus* | -0.0473 | 0.5269 | 0.016 | -, ** |
| 5 | *Saurida elongata* | -0.0290 | 0.4263 | 0.006 | -, n.s. |
| 6 | *Scolopsis taenioptera* | 0.0203 | 0.2221 | 0.002 | +, n.s. |
| 7 | Sciaenidae | -0.0594 | 0.5700 | 0.036 | -, ** |
| 8 | *Rastrelliger kanagurta* | -0.0983 | 0.6549 | 0.183 | -, *** |
| 9 | *Sepia aculeata* | -0.1017 | 0.6445 | 0.028 | -, n.s. |
| 10 | *Rastrelliger brachysoma* | 0.0943 | -0.3214 | 0.919 | +, n.s. |
| 11 | *Encrasicholina* spp. | 0.0683 | -0.1065 | 0.015 | +, n.s. |
| 12 | *Upeneus sulphureus* | -0.0806 | 0.9205 | 0.025 | -, * |
| 13 | Platycephalidae | -0.1348 | 1.3418 | 0.062 | -, *** |
| 14 | *Saurida undosquamis* | -0.0673 | 0.7578 | 0.028 | -, *** |
| 15 | *Nemipterus mesoprion* | -0.1025 | 1.0558 | 0.053 | -, *** |
| 16 | *Nemipterus peronii* | -0.0202 | 0.4845 | 0.002 | -, n.s. |
| 17 | *Nemipterus japonicus* | -0.0657 | 0.7656 | 0.024 | -, * |
| 18 | *Saurida isarankurai* | -0.1200 | 1.2020 | 0.075 | -, *** |
| 19 | *Atule mate* | -0.1642 | 1.6395 | 0.115 | -, *** |
| 20 | *Lagocephalus spadiceus* | -0.0865 | 0.8787 | 0.038 | -, * |
| 21 | *Siganus canaliculatus* | -0.1117 | 1.0659 | 0.133 | -, *** |
| 22 | *Sphyraena jello* | -0.0879 | 0.9350 | 0.053 | -, *** |
| 23 | *Megalaspis cordyla* | -0.1319 | 1.2658 | 0.114 | -, *** |
| 24 | *Parastromateus niger* | -0.0971 | 0.9634 | 0.069 | -, * |
| 25 | *Lutjanus lutjanus* | -0.0745 | 0.6082 | 0.205 | -, *** |

*a* is the density-dependent slope and *b* is the *y*-intercept of the virtual population dynamic model. ‘-’ denotes a negative relationship and ‘+’ a positive relationship between frequency at age *t* and survival rate of consequence age *t* + 1. Significance codes (linear regression): ‘\*\*\*’ $p < 0.001$ , ‘\*\*’ $p < 0.01$ , ‘\*’ $p < 0.05$ , ‘n.s.’ not significant

## Discussion

This study addressed the long-standing uncertainty surrounding the role of density-dependent mortality during early life stages in the tropical multispecies and multi-gear fisheries in the GoT. By integrating survey data with landing statistics from trawl fisheries, whose catches are dominated by TF mixed with juveniles, the results provide empirical evidence that density-dependent mortality is common among early life stages of fish in the GoT. Linking temporal variation in the abundance of individuals smaller than *L*_25_ to subsequent survival across multiple taxa revealed particularly strong compensatory signals among fast-growing and high-turnover taxa, whereas evidence for depensatory dynamics was weak or absent.

### Early-life density-dependent regulation in a tropical multispecies and multi-gear system

The growth parameters highlight pronounced variation in life-history strategies among taxa retained by trawlers in the GoT from 2016 to 2023, spanning small, fast-turnover pelagic species to larger, slower-growing demersal species (Table 2). This heterogeneity aligns with expectations for tropical trawl assemblages (Pauly and Christensen 1995) and supports a differential capacity for compensatory responses under exploitation. High growth performance (*ϕ*^′^) values for species such as *Sphyraena jello* and *Atule mate* indicate potentially rapid turnover and resilience, whereas lower *K* values for groups like the Sciaenidae imply slower recovery, reinforcing that juvenile harvesting is unlikely to exert uniform effects across species (Lorenzen 2008; Froese et al. 2014).

Density-dependent survival patterns further support theoretical expectations that compensatory mortality regulates pre-recruit populations (Jenkins et al. 1991; Myers and Cadigan 1993; Doherty et al. 2004; Ohlberger et al. 2014; Andersen et al. 2017; Houk et al. 2018; Lorenzen and Camp 2018; Zimmermann et al. 2018; Stige et al. 2019; Taylor et al. 2019; Houk et al. 2020). About 80% of the top 25 taxa exhibited negative survival–abundance slopes (Table 3), with several taxa showing strong and highly significant density dependence. These patterns indicate that compensatory mortality operated over a limited density range, within which a simple first-order response was sufficient to capture the dominant empirical biological signal, despite the potential for more complex nonlinear dynamics.

These results suggest that recruitment regulation may occur well before fisheries interception for many species, helping to explain why juvenile removal in multispecies tropical systems does not necessarily translate into reduced long-term productivity (Hazlerigg et al. 2012). Conversely, weak or non-significant density dependence observed in some taxa indicates that compensatory strength may vary among species or be masked by environmental variability, consistent with observations for other marine systems (Rose et al. 2001; Stige et al. 2019).

### Toward evidence-based management of TF under compensatory mortality

Although ecological resilience was evident in the GoT (Yuttharax et al. 2026), resiliency alone cannot substitute for management. Sustaining this system will require explicitly recognizing TF as a managed output, improving size-based governance, strengthening monitoring, and incorporating density-dependent processes into stock assessment frameworks. Such an approach would support Thailand in transitioning from incidental TF utilization toward evidence-based management that safeguards both ecological productivity and feed-supply resilience.

The findings of this study suggest that compensatory density-dependent mortality operates among many of the species commonly labeled TF. When compensatory dynamics exist within the array of TF, conventional population models assuming constant natural mortality (Hilborn and Walters 1992; Quinn and Deriso 1999; Brodziak et al. 2011; Haddon 2011) are likely to overestimate the impacts of pre-recruitment fishing (Fig. 2), as increases in juvenile mortality are partially offset by density-dependent survival. Consequently, the marginal effect of juvenile removal on long-term productivity may be more limited than usually assumed.

**Fig. 2:**
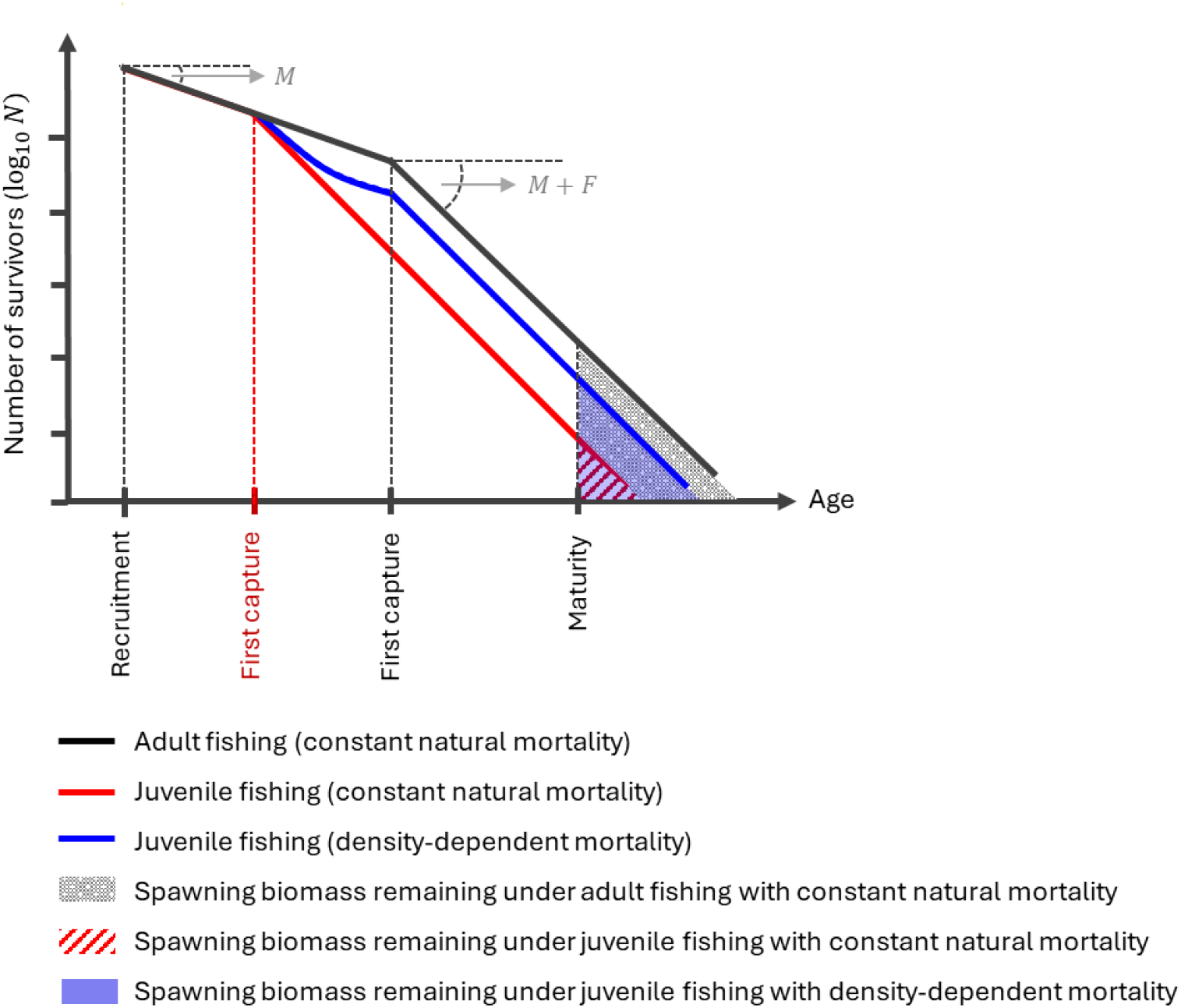
Conceptual illustration of density-dependent mortality and its implications for stock assessment

Within this context, the spawning potential ratio (SPR) framework (Mace 1994; Gabriel and Mace 1999) provides a useful conceptual reference. SPR theory indicates that maintaining spawning biomass at approximately 30%–40% of unfished levels is generally sufficient to sustain reproductive capacity, and that further increases beyond this threshold do not necessarily yield proportional gains in sustainability. When compensatory density-dependent mortality is present, this framework remains consistent with the expectation that increases in spawning biomass above critical thresholds may produce diminishing returns.

However, although the existence of compensatory density-dependent mortality is supported, its magnitude among TF species in the GoT cannot yet be quantified with sufficient precision to establish operational reference points. Future efforts should therefore focus on refining sampling designs and on empirical estimations of early-life density dependence. Improved parameterization will be essential for incorporating compensatory processes into assessment models and for advancing evidence-based management of TF resources. These recommendations extend beyond Thailand and are equally relevant to tropical fisheries worldwide, particularly in regions where TF are intensively utilized for non-food purposes.

Taken together, these findings indicate that TF assemblages possess inherent demographic buffering. However, the lack of a compensatory signal in some slower-growing taxa underscores that resilience is not universal. Species-specific differences in growth and compensatory capacity, therefore, warrant attention in management considerations, particularly where trawl fisheries operate non-selectively. This study provides a better understanding of how abundance interacts with density-regulated survival, by providing a biological basis for evaluating the implications of juvenile extraction. The findings should inform more nuanced perspectives on balanced harvest strategies in tropical multispecies and multi-gear fisheries.

## Acknowledgements

The following persons coordinated and delivered the data to the researchers: Napat Mahasawat, Jidapa Taweteekul, Sakol Pheaphabrattana, Prompan Hiranmongkolrat, and Weerapol Thitipongtrakul for the map preparation (Department of Fisheries, Thailand). Generative AI tools were initially used to assist with writing in English and literature searches. Cynthia Kulongowski, a subject-expert editor with Edanz (https://jp.edanz.com/ac), edited the language of a draft of this manuscript.

## Author contributions

JY conducted the data analysis, visualization, and manuscript writing. CMF contributed to the development of scripts for the growth-parameter analysis through comparative evaluation. PN and MK reviewed the manuscript and guided the interpretation of the results in the context of Thailand’s fisheries resources and marine ecosystem. MTF verified the analytical methods, revised the manuscript, and supervised the findings of this work. All authors contributed to the discussion of the results.

## Data availability

All the sources of data used in this study are cited within the text. Only the commercial trawl-landing survey data are subject to permission from the Department of Fisheries (DoF), Thailand. However, because of security concerns related to potential cyber threats, the online data of the DoF website may occasionally limit access from outside the country (or for some countries); therefore, contact with the DoF is generally needed to access those data.

## References

Andersen KH, Jacobsen NS, Jansen T, Beyer JE (2017) When in life does density dependence occur in fish populations? Fish Fish 18:656–667. 10.1111/faf.12195

Beverton RJH, Holt SJ (1957) On the dynamics of exploited fish populations. Fishery investigations series II, Volume 19. Ministry of Agriculture, Fisheries and Food, London

Broadhurst MK, Suuronen P, Hulme A (2006) Estimating collateral mortality from towed fishing gear: collateral mortality from towed gear. Fish and Fisheries 7(3):180–218. 10.1111/j.1467-2979.2006.00213.x

Brodziak JK, Ianelli JN, Lorenzen K, Methot RD (Eds.) (2011) Estimating natural mortality in stock assessment applications. US Department of Commerce, National Oceanic and Atmospheric Administration, National Marine Fisheries Service.

Carpenter KE, Niem VH (1998a) FAO species identification guide for fishery purposes. The living marine resources of the Western Central Pacific Vol 1. Seaweeds, corals, bivalves and gastropods. FAO, Rome

Carpenter KE, Niem VH (1998b) FAO species identification guide for fishery purposes. The living marine resources of the Western Central Pacific Vol 2. Cephalopods, crustaceans, holothurians and sharks. FAO, Rome

Carpenter KE, Niem VH (1999a) FAO species identification guide for fishery purposes. The living marine resources of the Western Central Pacific Vol 3. Batoid fishes, chiamaeras and bony fishes Part 1 (Elopidae to Linophrynidae). FAO, Rome

Carpenter KE, Niem VH (1999b) FAO species identification guide for fishery purposes. The living marine resources of the Western Central Pacific Vol 4. Bony fishes Part 2 (Mugilidae to Carangidae). FAO, Rome

Carpenter KE, Niem VH (2001a) FAO species identification guide for fishery purposes. The living marine resources of the Western Central Pacific Vol 5. Bony fishes Part 3 (Menidae to Pomacentridae). FAO, Rome

Carpenter KE, Niem VH (2001b) FAO species identification guide for fishery purposes. The living marine resources of the Western Central Pacific Vol 6. Bony fishes Part 4 (Labridae to Latimeriidae), estuarine crocodiles, sea turtles, sea snakes and marine mammals. FAO, Rome

Christensen V (1996) Managing fisheries involving predator and prey species. Rev Fish Biol Fish 6:417–442. 10.1007/BF00164324

Doherty PJ, Dufour V, Galzin R, Hixon MA, Meekan MG, Planes S (2004) High mortality during settlement is a population bottleneck for a tropical surgeonfish. Ecology 85(9):2422–2428. 10.1890/04-0366

FAO (2011) International guidelines on bycatch management and reduction of discards. Food and Agriculture Organization of the United Nations, Rome

FAO (2022) The state of world fisheries and aquaculture – Towards blue transformation. FAO, Rome. 10.4060/cc0461en

FAO (2023) The status of marine fishery stock assessments in the Asian region and the potential for a network of practitioners. FAO, Bangkok

Froese R, Thorson JT, Reyes Jr RB (2014) A Bayesian approach for estimating length–weight relationships in fishes. Journal of Applied Ichthyology 30(1):78–85. 10.1111/jai.12299

Funge-Smith S, Lindebo E, Staples D (2005) Asian fisheries today: the production and use of low value/trash fish from marine fisheries in the Asia-Pacific region. RAP Publication 2005/16. https://openknowledge.fao.org/server/api/core/bitstreams/9f0e57bd-b244-4629-836e-e8aaaaf76bf7/content. Accessed 10 Oct 2025

Gabriel WL, Mace PM (1999) A review of biological reference points in the context of the precautionary approach. In proceedings of the fifth national NMFS stock assessment workshop providing scientific advice to implement the precautionary approach under the Magnuson-Stevens fishery conservation and management act; NOAA Technical Memorandum NMFS-F/SPO-4, USA

Gilman E, Suuronen P, Hall M, Kennelly S (2013) Causes and methods to estimate cryptic sources of fishing mortality. Journal of Fish Biology. 10.1111/jfb.12148

Haddon M (2011) Modelling and quantitative methods in fisheries. Chapman and Hall CRC Press, London, 405

Hazlerigg CRE, Lorenzen K, Thorbek P, Wheeler JR, Tyler CR (2012) Density-dependent processes in the life history of fishes: Evidence from laboratory populations of zebrafish Danio rerio. PLOS ONE 7(5):e37550. 10.1371/journal.pone.0037550

He P (2015) Systematic research to reduce unintentional fishing-related mortality: example of the Gulf of Maine northern shrimp trawl fishery. In Kruse G, An H, DiCosimo J, Eischens C, Gislason G, eds. Fisheries bycatch: Global issues and creative solutions, p. Alaska Sea Grant, University of Alaska Fairbanks.

Hilborn R, Walters CJ (1992) Quantitative fisheries stock assessment: choice, dynamics and uncertainty. Chapman and Hall, New York

Houk P, Cuetos-Bueno J, Tibbatts B, Gutierrez J (2018) Variable density dependence and the restructuring of coral-reef fisheries across 25 years of exploitation. Sci Rep 8:5725. 10.1038/s41598-018-23971-6

Houk P, Lemer S, Hernandez-Ortiz D, Cuetos-Bueno J (2020) Phylogenies predict compensatory density dependence in coral-reef fisheries. Authorea. 10.22541/au.159657670.03126007

Jenkins GP, Young JW, Davis TLO (1991) Density dependence of larval growth of a marine fish, the southern bluefin tuna, Thunnus maccoyii. Canadian Journal of Fisheries and Aquatic Sciences 48(8):1358–1363. 10.1139/f91-162

Kelleher K (2005) Discards in the world’s marine fisheries. An update. FAO, Rome

Kulanujaree N, Salin KR, Noranarttragoon P, Yakupitiyage A (2020) The transition from unregulated to regulated fishing in Thailand. Sustain. 10.3390/su12145841

Longhurst A, Pauly D (1987) Ecology of tropical oceans. Academic Press, San Diego

Lorenzen K (2000) Allometry of natural mortality as a basis for assessing optimal release size in fish stocking programmes. Can J Fish Aquat Sci 57(12):2374–2381. 10.1139/f00-215

Lorenzen K (2008) Fish population regulation beyond “stock and recruitment”: the role of density-dependent growth in the recruited stock. Bulletin of Marine Science 83(1):181–196

Lorenzen K, Camp EV (2018) Density-dependence in the life history of fishes: when is a fish recruited? Fisheries Research 217:5–10. 10.1016/j.fishres.2018.09.024

Mace PM (1994) Relationships between common biological reference points used as thresholds and targets of fisheries management strategies. Can J Fish Aquat Sci 51(1):110–122. 10.1139/f94-013

Main J, Sangster G (1990) An assessment of the scale damage to and survival rates of young fish escaping from the cod-end of a demersal trawl. Scottish Fisheries Research Report (46):28

Matsuishi TF (2025) Status of Southeast Asian fisheries: distinctive characteristics and pathways to sustainable fisheries. Fish Sci. 10.1007/s12562-025-01854-w

Mildenberger TK, Taylor MH, Wolff M (2017) TropFishR: an R package for fisheries analysis with length–frequency data. Methods in Ecology and Evolution 8(11):1520–1527. 10.1111/2041-210X.12791

Myers RA, Cadigan NG (1993) Density-dependent juvenile mortality in marine demersal fish. Can J Fish Aquat Sci 50(8):1576–1590

Noranarttragoon P, Koolkalaya S, Thitipongtrakul W, Avakul P, Phoonsawat R, Jutagate T (2023) Trawl fisheries in the Gulf of Thailand: vulnerability assessment and trend analysis of the fish landings. Fishes 8. 10.3390/fishes8040177

Ohlberger J, Rogers LA, Stenseth NC (2014) Stochasticity and determinism: how density-independent and density-dependent processes affect population variability. PLOS ONE 99(6):e98940. 10.1371/journal.pone.0098940

Pauly D (1980) A selection of simple methods for the assessment of tropical fish stocks. FAO Fisheries Circular No 729, FAO, Rome

Pauly D, Christensen V (1995) Primary production required to sustain global fisheries. Nature 374: 255–257. 10.1038/374255a0

Pauly D, Christensen V, Dalsgaard J, Froese R, Torres F (1998) Fishing down marine food webs. Science (80)279:860–863. 10.1126/science.279.5352.860

Pauly D, Munro JL (1984). Once more on the comparison of growth in fish and invertebrates. Fishbyte 2: 21

Pérez Roda MA, Gilman E, Huntington T, Kennelly SJ, Suuronen P, Chaloupka M, Medley P (2019) A third assessment of global marine fisheries discards. Rome

Pope J (1979) Stock assessment in multispecies fisheries, with special reference to the trawl fishery in the Gulf of Thailand. South China Sea Fisheries Development and Coordinating Programme, FAO, Manila

Quinn TJ, Deriso RB (1999) Quantitative fish dynamics. Oxford University Press.

Ricker WE (1954) Stock and recruitment. Journal of the Fisheries Research Board of Canada 11:559–623. 10.1139/f54-039

Rose KA, Cowan Jr JH, Winemiller KO, Myers RA, and Hilborn R (2001) Compensatory density dependence in fish populations: importance, controversy, understanding and prognosis. Fish and Fisheries 2(4):293–327

Schwamborn R, Freitas MO, Moura RL, Aschenbrenner A (2023) Comparing the accuracy and precision of novel bootstrapped length-frequency and length-at-age (otolith) analyses, with a case study of lane snapper (Lutjanus synagris) in the SW Atlantic. Fisheries Research 264:106735. 10.1016/j.fishres.2023.106735

Schwamborn R, Mildenberger TK, Taylor MH (2019) Assessing sources of uncertainty in length-based estimates of body growth in populations of fishes and macroinvertebrates with bootstrapped ELEFAN. Ecological Modelling 393:37–51. 10.1016/j.ecolmodel.2018.12.001

Sherman K, Hempel G (2009) The Unep Large Marine Ecosystems Report: A perspective on changing conditions in LMEs of the world’s regional seas. Nairobi, Kenya

Sparre P, Venema SC (1998) Introduction to tropical fish stock assessment. Part 1: Manual. FAO Fisheries Technical Paper No. 306.1, Rev. 2. FAO, Rome

Staples D, Funge-Smith S (2005) Prized commodity: low-value/trash fish from marine fisheries in the Asia-Pacific region. Fish People 3:1–15

Stige LC, Rogers LA, Neuheimer AB, Hunsicker ME, Yaragina NA, Ottersen G, Ciannelli L, Langangen O, Durant JM (2019) Density- and size-dependent mortality in fish early life stages. Fish and Fisheries 20:962–976. 10.1111/faf.12391

Suuronen P (2005) Mortality of fish escaping trawl gears. FAO Fisheries Technical Paper (478). FAO, Rome

Suuronen P, Sardà F (2007) The role of technical measures in European fisheries management and how to make them work better. ICES Journal of Marine Science 64:751–756

Taufani WT, Matsuishi TF (2025a) Precision and accuracy in estimating biological reference points using length-based fisheries assessment: a bootstrapping approach. Fisheries Science 91:1175–1187. 10.1007/s12562-025-01921-2

Taufani WT, Matsuishi TF (2025b) Precision of estimated growth parameters of yellowfin tuna (Thunnus albacares) from length-frequency data estimated by bootstrapping. Fisheries Management and Ecology 32:50–58. 10.1111/fme.12781

Taylor BM, Choat JH, DeMartini EE, Hoey AS, Marshell A, Priest MA, Rhodes KL, Meekan MG (2019) Demographic plasticity facilitates ecological and economic resilience in a commercially important reef fish. J Anim Ecol 88:1888–1900. 10.1111/1365-2656.13095

Yuttharax J, Noranarttragoon P, Thitipongtrakul W, Matsuishi TF (2026) Ecological reality of trash fish catches from trawlers under tropical multispecies and multi-gear fisheries in the Gulf of Thailand. Fish Sci. 10.1007/s12562-025-01952-9

Zimmermann F, Ricard D, Heino M (2018) Density regulation in Northeast Arctic fish populations: density dependence is stronger in recruitment than in somatic growth. Journal of Animal Ecology 87:672–681. 10.1111/1365-2656.12800

